# A DNA condensation code for linker histones

**DOI:** 10.1101/2023.11.20.567813

**Authors:** Matthew Watson, Dilyara Sabirova, Megan C. Hardy, Yuming Pan, Henry Yates, Charlotte J. Wright, W. H. Chan, Ebru Destan, Katherine Stott

## Abstract

Linker histones play an essential role in chromatin packaging by facilitating compaction of the 11-nm fibre of nucleosomal “beads on a string”. The result is a heterogeneous condensed state with local properties that range from dynamic, irregular and liquid-like, to stable and regular structures (the 30-nm fibre), which in turn impact chromatin-dependent activities at a fundamental level. The properties of the condensed state depend on the type of linker histone, particularly on the highly disordered C-terminal tail, which is the most variable region of the protein, both between species, and within the various subtypes and cell-type specific variants of a given organism. We have developed an *in-vitro* model system comprising linker histone tail and linker DNA, which although very minimal, displays surprisingly complex behaviour, and is sufficient to model the known states of linker-histone-condensed chromatin: disordered “fuzzy” complexes (“open” chromatin), dense liquid-like assemblies (dynamic condensates) and higher-order structures (organised 30-nm fibres). A crucial advantage of such a simple model is that it allows the study of the various condensed states by NMR, CD and scattering methods. Moreover, it allows capture of the thermodynamics underpinning the transitions between states through calorimetry. We have leveraged this to rationalise the distinct condensing properties of linker histone subtypes and variants across species that are encoded by the amino acid content of their C-terminal tails. Three properties emerge as key to defining the condensed state: charge density, lysine/arginine ratio, and proline-free regions, and we evaluate each separately using a strategic mutagenesis approach.

## Introduction

Chromatin – the complex of genomic DNA and proteins in eukaryotes – is packaged in stages (1). The basic unit is the nucleosome (2), in which the DNA performs almost two full turns around an octamer of core histones. Nucleosomes form arrays of “beads on a string” that make up an 11 nm fibre. The next stage, in which linker histones bring about further condensation, is more enigmatic, despite it spanning a length scale that is critical for the control of key transactions such as gene expression, replication and repair (3). The orderly 30-nm fibre originally proposed (4) has been detected *in vivo* (5, 6), but only in avian erythrocytes, that are terminally-differentiated and transcriptionally-silent. More recently, cryo-ET and live-cell imaging have revealed that the 11 nm fibre compacts via irregular folding (7) and is dynamic (8, 9), specifically that clusters of nucleosomes show coherent motions suggestive of liquid flow (10, 11). It is likely that, at the local scale, linker-histone condensed chromatin can adopt a virtual continuum of states between solid and liquid, and that these impact the accessibility of the underlying DNA: a liquid-like state is a compelling means by which chromatin could respond quickly to environmental stimuli, allowing transactions such as transcription, replication and repair. Conversely, unfolding and refolding a fibre-like architecture would be much less efficient.

The growing appreciation of dynamics in chromatin packaging by ‘top-down’ imaging has paralleled developments by us and others in our ‘bottom-up’ understanding of the linker histone proteins at the molecular level. Linker histones consist of a winged-helix domain (∼80 residues) flanked by short N-terminal and long C-terminal disordered regions (25-30 and ∼100 residues, respectively). The globular domain locates H1 at the nucleosome dyad (12), and along with the first 13-residues of the C-terminal domain, stabilises the entering and exiting DNA strands in a stem arrangement (13). The rest of the C-terminal domain binds the linker DNA between nucleosomes to facilitate their closer approach, but is not well defined by cryo-EM (14) and invisible by cryo-ET (7) due to dynamics (15). Previously, we studied the isolated C-terminal domain (‘CH1’) in order to reveal its structure and condensation behaviour with DNA by NMR spectroscopy and isothermal titration calorimetry (ITC) (16). This model system of CH1 and linker DNA, although very minimal, displays surprisingly complex behaviour, forming three distinct states depending on the conditions: disordered complexes, phase separated droplets and higher-order structures, which appear to have some correspondence to the states of H1-bound chromatin (“open” 11 nm fibres, dense liquid-like assemblies and organised 30-nm fibres, respectively) (17) (Fig. 1). The sufficiency of CH1 for DNA condensation is consistent with its requirement for high affinity chromatin binding (18), its essential role in condensation of native chromatin beyond the 11 nm fibre (19), and as the main driver for phase separation and decreased dynamics in a subsequent study of *in vitro* reconstituted 12-nucleosome arrays (20).

**Figure 1.**
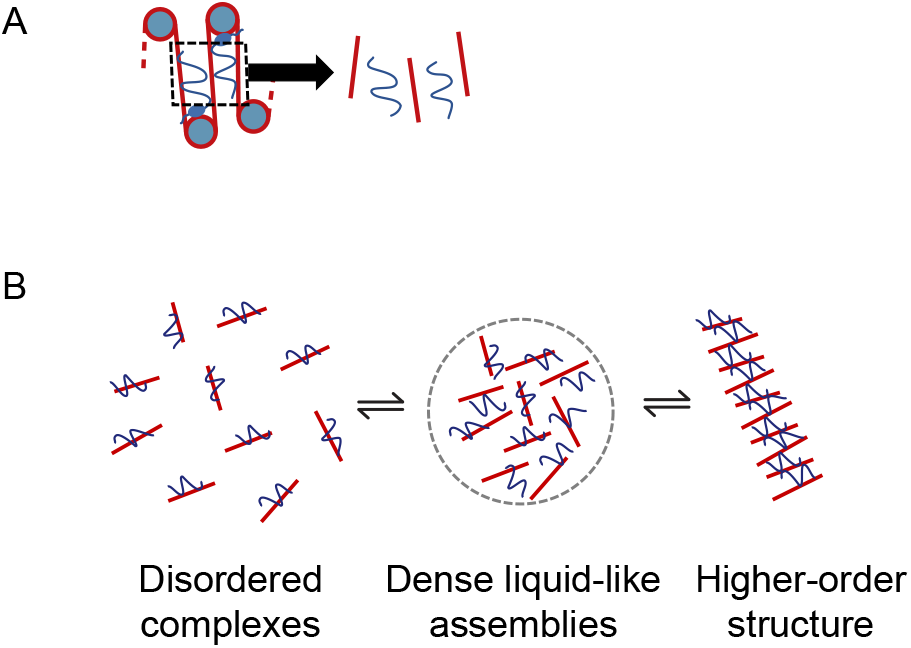
A model encompassing the possible H1-condensed chromatin states. (A) A minimal model for the study of chromatin condensation by linker histones consists of internucleosomal linker DNA and the C-terminal tail of H1 (CH1) (16). (B) The model system forms disordered complexes, phase separated droplets, and higher-order structures (a liquid crystalline phase), depending on the conditions.

All proteins that condense DNA are rich in lysine, arginine, or both. The linker histone tails are essentially co-polymers of lysine, alanine and proline, with variable amounts of arginine, depending on the subtype/variant. At a phenomenological level, arginine has been observed to generate more dramatic DNA condensation than lysine. This is due to the distinct chemistry of their side-chains: the amino group of lysine makes weak ion-pairing interactions with the DNA phosphates, while the guanidino group of arginine has the propensity to make additional π-stacking and bidentate interactions, and hydrogen bonds (21–24). It follows that the amino acid content of linker histone tails is likely to code for a range of condensed chromatin ‘baseline’ states that differ in their degree of compaction and other material properties. Eukaryotes make several different H1 subtypes (for example there are 11 in humans and mice), which can differ in their expression across cell types and their localisation (25–27). Some are expressed in a replication-dependent manner, while others occur exclusively in germ cells. Further, several variant linker histones (e.g. H5 in *Gallus gallus* (28)) are associated with terminally-differentiated cells. The winged-helix globular domains are highly conserved across linker histones, albeit with minor differences (five residues) that determine the exact positioning of the globular domain relative to the nucleosome dyad (29). Therefore, differences in the intrinsic condensing properties are likely to arise from sequence and content variation in the tail (this is already known to affect the orientation of the linkers directly flanking the nucleosome (30)), or from distinct post-translational modifications (31). In this study, we explicitly address the question of how the natural variation in linker histone sequences directly affects DNA condensation by comparison of the structure, thermodynamics and condensation propensities of tails with a range of potentially relevant properties spanning different lysine/arginine ratios, positive charge densities, hydrophobicity and proline content and spacing.

## Results

### Linker histone H1 tails are universally well-mixed polycationic electrolytes

In order to understand the amino acid sequence space of the known linker histone H1 tails, we curated a database free from erroneous inclusions and annotation errors. An initial set of 245 sequences was revealed from a search for “Histone H1” in UniProtKB/SwissProt after exclusion of sequences from bacteria and viruses. Further manual filtering to remove hypothetical proteins, protamines, and sequences with no globular domain or significant tails reduced the number to 94. The sequences were aligned on the globular domain in order to extract the C-terminal tails (Supp. Fig. S1), which were then analysed for content, charge, and distribution of charge and proline (Fig. 2). The tails are dominated by lysine (mean 34.6%, σ 4.2%) and alanine (mean 24.0%, σ 4.1%), followed by proline (mean 10.6%, σ 1.7%) (Fig. 2A). They cluster on a Das-Pappu phase diagram of states (32, 33) around an average *f*_+_, *f*_–_ coordinate of (0.38, 0.03), indicating a very high fraction of positively charged residues and relatively few negatively charged residues (Fig. 2B). Charge patterning was assessed by comparing *κ* parameters across the set (Fig. 2C). *κ* spans values between 0 (well-mixed/fully alternating) to 1 (fully segregated); the distribution of *κ* in CH1s is concentrated around 0.13, indicating that the charge is well mixed in all linker histones of the set. The distribution of *Ω* (which includes proline as well as charge) showed an even greater level of mixing, centred around 0.06. Taken together, the set of linker histone tails is one of universally well-mixed polycationic electrolytes that are conserved in coarse-grained features (Supp. Fig. S1).

**Figure 2.**
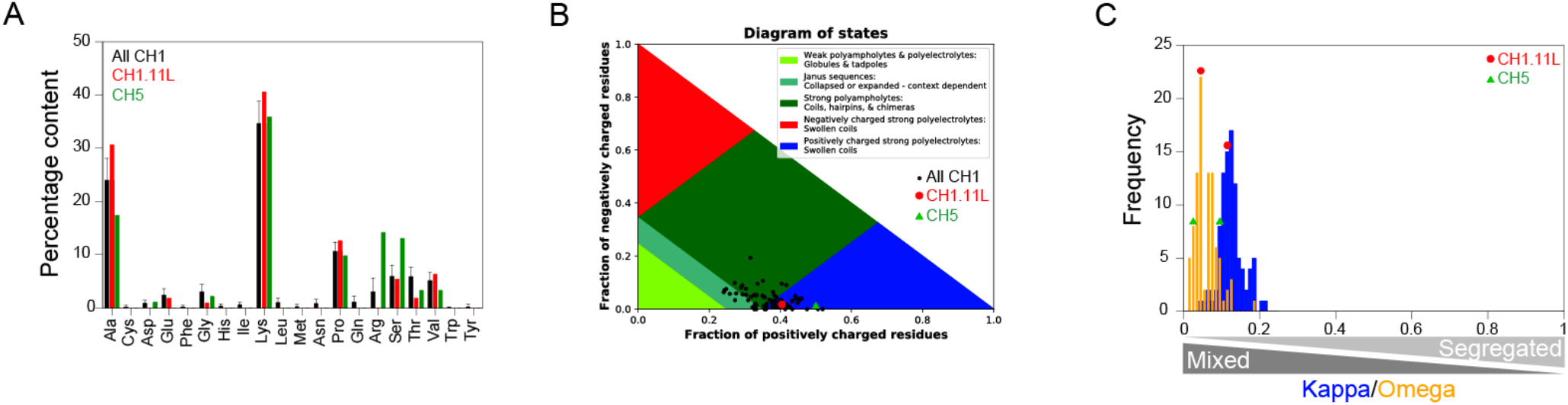
Sequence content and physical properties of H1 C-terminal tails (CH1s). (A) Amino acid residue content of CH1s by percentage. (B) Curated set of 94 CH1s displayed on a phase diagram of IDP states (32, 33) (*Gallus gallus* isoform H1.11L and variant H5 are shown as a red circle and a green triangle, respectively). (C) Distributions of *κ* (charge patterning; blue) and *Ω* (patterning of charged and proline residues; orange) where 0 = well mixed, 1 = fully segregated.

### Sampling sequence space *via* mutagenesis

We have studied H1.11L from *Gallus gallus* experimentally as a model linker histone for many years (16, 34). It is typical of the somatic isoforms in chicken as well as the larger set collated here, being 41% lysine, which is well distributed (*κ* = 0.1) (Fig. 2A). The remaining residues are mainly alanine (31%) and proline (13%), which is also well distributed (*Ω* = 0.05). The most hydrophobic residue is valine, of which there are seven. The positions of both proline and valine are well conserved across the six somatic chicken isoforms (Supp. Fig. S2A), although the valine is not always present (two of the isoforms have only two valines). Chicken H1s contain little, if any, arginine. However, a natural variant found in avian erythrocytes, H5 (28), contains 13 arginine residues in its C-terminal tail (Supp. Fig. S2B), and is also more charge dense, being 50% lysine/arginine (Fig. 2A&B). Notably, this variant forms detectable 30-nm fibres *in vivo* (6).

The natural variation in linker histone tail sequences exemplified by the chicken isoforms and variants spans many properties that could potentially impact DNA condensation: lysine/arginine ratio, positive charge density and hydrophobicity. The high proline content is also intriguing. These unusual proteins are challenging to express and purify, therefore a minimal set of mutants was designed that could reveal the impact of the amino acid content in as systematic a manner as possible (Fig. 3). The set consists of the C-terminal tails of H1.11L and H5 (denoted CH1 and CH5), two different valine knockouts (CH1_VA_ and CH1_VT_), and a proline knockout (CH1_PA_). For the valine knockouts, a substitution by threonine was tested as well as by alanine since threonine has a similar bulk to valine, but higher polarity than alanine. Further, in order to deconvolute the effect of arginine content and charge density, two additional proteins were generated: CH5_K_, in which the 13 arginine residues were mutated to lysine (thus maintaining the charge density of CH5 while removing the arginine), and CH1_R_, in which 13 of the lysines in CH1 were mutated to arginine in roughly the same patterning as for the arginines in CH5 (thus maintaining the charge density of CH1 but with an arginine:lysine ratio of 0.4, as for CH5).

**Figure 3.**
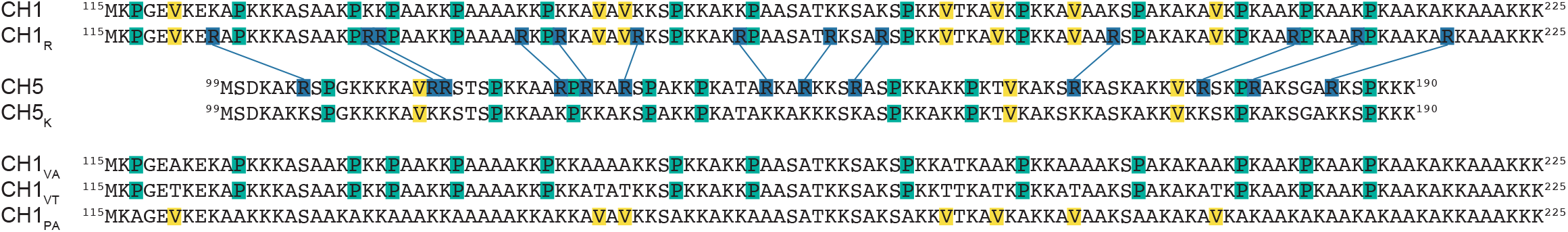
Mutant sequences. CH1 and CH5 are the wild-type sequences of the C-terminal tails of H1.11L and H5 from *Gallus gallus*. For the rationale behind the design of CH1_R_, CH5_K_, CH1_VA_, CH1_VT_ and CH1_PA_ see the main text. Arginine, proline and valine are highlighted in dark blue, green and yellow respectively.

### DNA condensation is driven by both positive charge density and arginine content

The condensation of 20bp dsDNA by the proteins and mutants was assessed by A_340_ (turbidity), by mixing at 1:1 molar stoichiometry at the concentrations shown, followed by addition of salt, to produce phase diagrams. Comparison of CH1 and CH5 shows that condensate formation by CH5 is more robust at lower concentrations and persists at higher ionic strengths (Fig. 4A). The CH1_R_ and CH5_K_ mutants were intermediate in their condensing properties, and broadly similar, demonstrating that the condensing power of CH5 derives from the additive effects of (i) its 13 arginines in place of lysine, and (ii) its 10% higher charge density.

**Figure 4.**
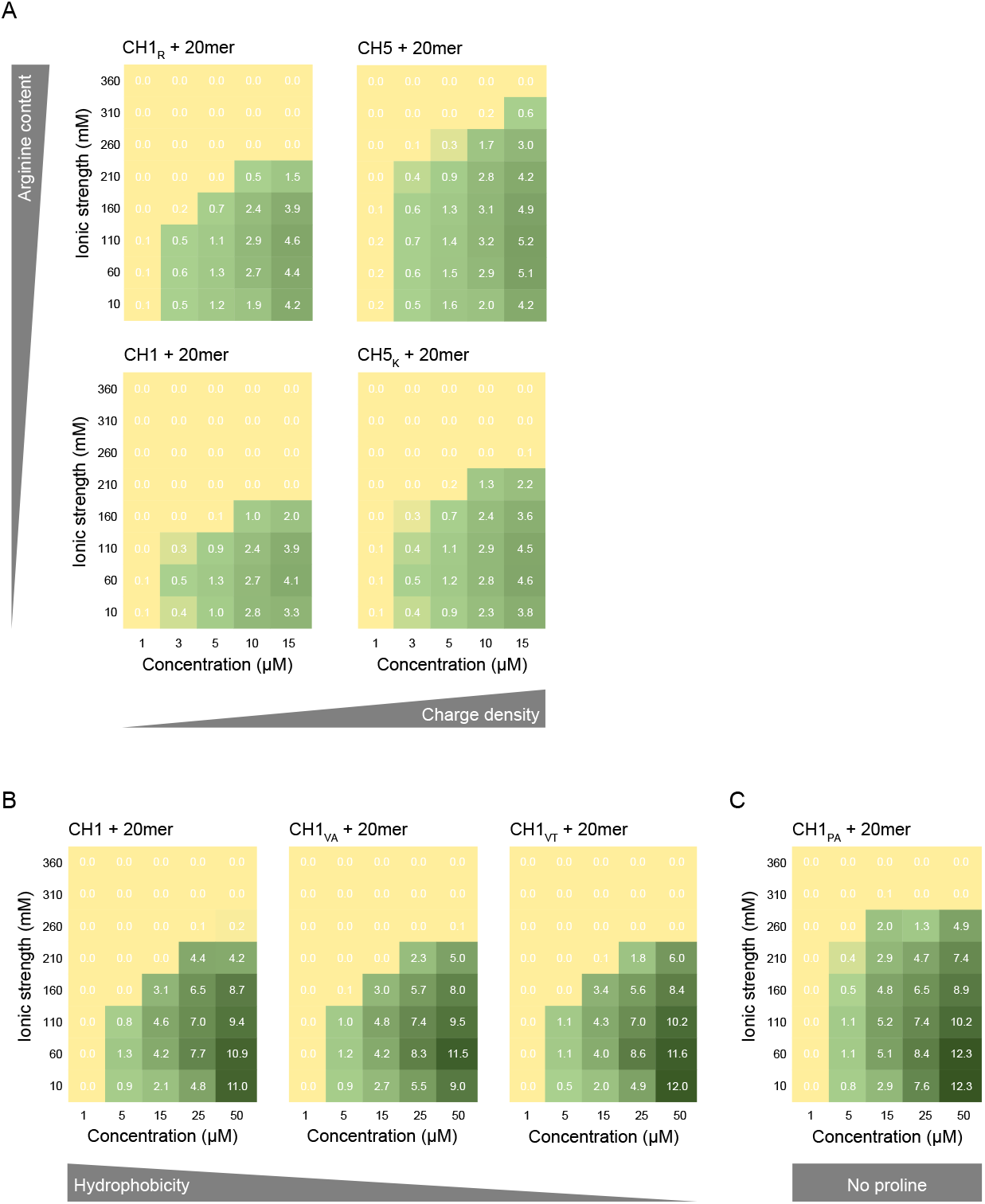
Phase diagrams with DNA vs concentration and ionic strength. (A) Arginine and charge density series. (B) Hydrophobicity series. (C) Proline knockout. Turbidity (A_340_) of the 1:1 complex is shown, in numbers as well as a yellow-to-green colour scale.

### Proline negatively regulates DNA condensation

The phase diagrams of CH1 and the less hydrophobic CH1_VA_ and CH1_VT_ were very similar (Fig. 4B). Therefore, the seven valines in CH1 do not significantly affect the DNA condensing power of the protein, suggesting that it is entirely driven by electrostatics. In order to verify this, the condensates were challenged with 1,6- hexanediol, a known disrupter of hydrophobically-driven condensate formation (35). Very little change was seen, for example, an addition of 10% 1,6-hexanediol by volume – the highest amount commonly used – to a CH1:20mer condensate lowered the A_340_ from 3.88 to 3.67, a reduction of only 5% (not shown). In contrast to the minimal effect of valine mutagenesis, CH1_PA_ had a dramatic effect on condensation, being more robust at lower concentrations and persisting to higher ionic strength (Fig. 4C). This was explored in further experiments.

### Robust condensation is underpinned by enthalpic contributions

The thermodynamics driving condensation of 20bp dsDNA by the natural proteins and mutants was investigated by ITC. This method, when coupled with a parallel titration followed by turbidity, reveals the heats associated with DNA binding and phase separation and is able to separate the contributions to some degree (16). The isotherms for CH1 and CH5 showed endothermic (positive) heats, albeit of different magnitudes (Fig. 5A). Condensation is therefore an entropy driven process, likely through counter-ion release (36). The isotherms display varying degrees of biphasic behavior due to the onset of phase separation, which results in a characteristic ‘hump’. This is confirmed by comparison with the turbidity titrations (allowing for the fact that turbidity is a cumulative measurement while ITC gives the heat per injection: the peak in the turbidity measurement therefore corresponds to the maximum downward gradient in the isotherm at *N* ∼ 1, equivalent to the inflexion point were it a simple sigmoid). The initial injections where little or no phase separation is occurring therefore reflect ion pairing (protein-DNA binding), and the later injections the process of ion pairing due to binding and phase separation. This phenomenon has been discussed previously (16, 37).

**Figure 5.**
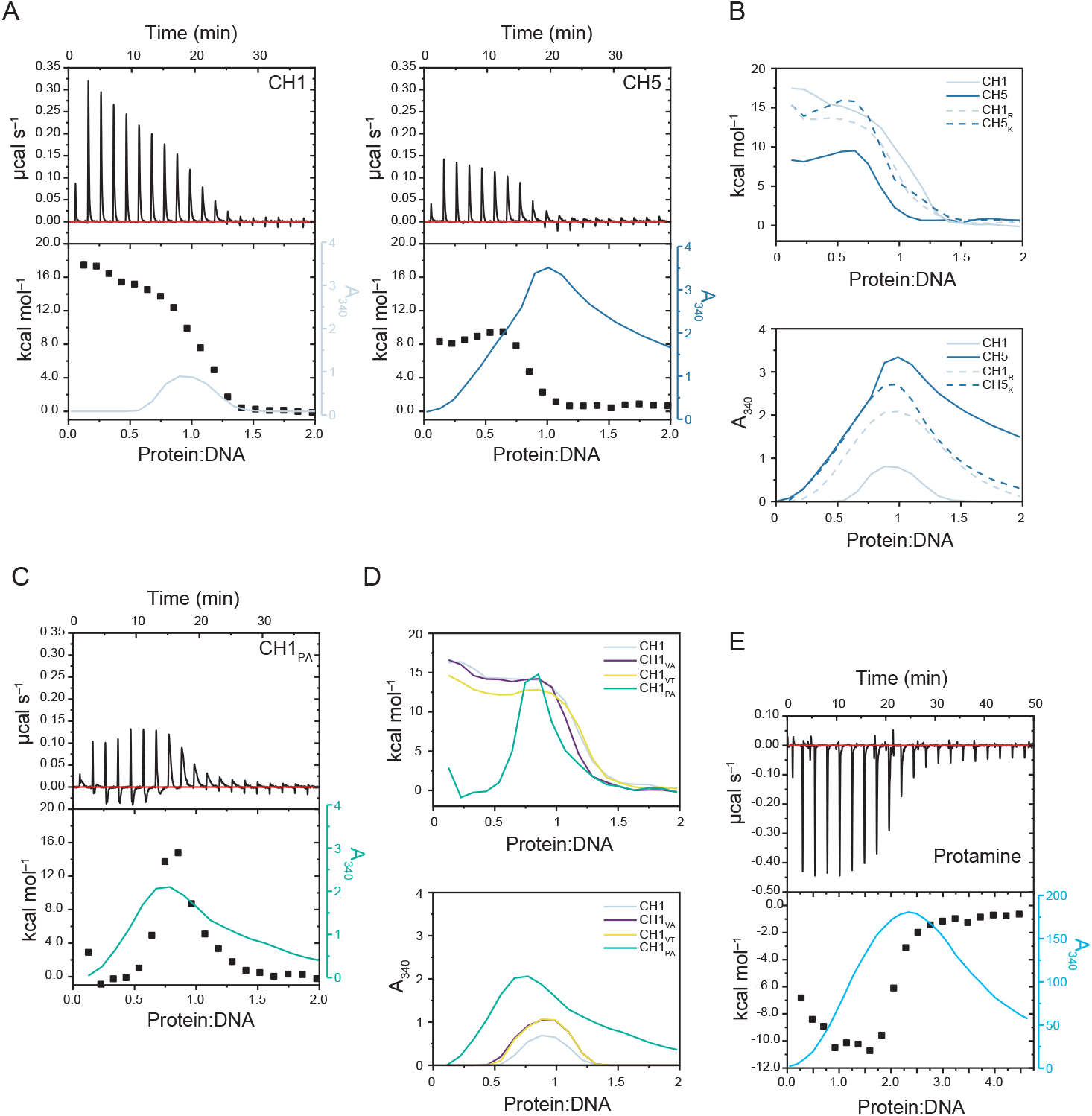
Isothermal titration calorimetry. Titrations of protein (as indicated) into 20 bp DNA, alongside A_340_ to identify phase separation events, in buffer containing 150 mM NaCl (total *I* = 160 mM). (A) Raw data, isotherms and A_340_ for CH1 and CH5. (B) Isotherms (above) and A_340_ (below) for the complete arginine and charge density series. (C) Raw data, isotherms and A_340_ for the proline knockout. (D) Isotherms (above) and A_340_ (below) for the hydrophobicity and proline knockouts. (E) Raw data, isotherms and A_340_ for Protamine titrated into 16 bp DNA. Complete dataset in Supp. Fig. S3.

The isotherms for CH1_R_ and CH5_K_ showed intermediate features (Supp. Fig. S3). Comparison of integrated and normalized heats of injection for CH1, CH5, CH1_R_ and CH5_K_ (Fig. 5B) reveals that CH5 gives the lowest endothermic heats (*y*-intercept +8 kcal/mol) and CH1 the highest (+17 kcal/mol). The mutants CH1_R_ and CH5_K_ are in between (+15 kcal/mol). Following this initial phase, the ‘hump’ is most pronounced for CH5 and CH5_K_, but more subtle for CH1 and CH1_R_. The turbidities (Fig. 5B) show that phase separation for CH5 has the earliest onset, reaches the highest values, and persists to the highest molar ratio, i.e. resists re-entrant behavior through over-charging. CH1 phase separates to a limited degree around *N* = 1, which is consistent with the turbidity under the same conditions (10 μM concentration, *I* = 160 mM (Fig. 4A)) and the mutants are in between.

The lower hydrophobicity mutants, CH1_VA_ and CH1_VT_, showed similar thermodynamic signatures to CH1 (Supp. Fig. S3), although CH1_VT_ had slightly lower heat and higher *N*, and both mutants had slightly higher turbidity. However, CH1_PA_ was profoundly different (Fig. 5C). Around the onset of turbidity, which occurs almost immediately as for CH5, the isotherm became more complex, the peaks showing both endothermic (positive) and exothermic (negative) features. Despite this complexity, it was possible to link the observed thermodynamic events with phase separation by comparison with the turbidity and the other isotherms (Fig. 5D). The onset of phase separation is much earlier for CH1_PA_ than for CH1, correlating with exothermic signals in the first 7-8 injections. The exothermic signals were broad, in line with previous observations that signals with slow relaxations are highly characteristic of phase separation (37). Superimposed on these are the sharper endothermic peaks that are characteristic of ion pairing. At the peak in turbidity, which is earlier for CH1_PA_ at *N* ∼ 0.8, the broad signals change sign, consistent with progressive dissolution of the condensate through over-charging of the system with protein. These features are indicative that phase separation for CH1_PA_ is enthalpically driven. Overall therefore, it appears that robust condensation correlates with a reduction in endothermic heats for CH1_R_, CH5_K_ and CH5, switching to exothermic for CH1_PA_. This trend was further explored by testing a highly charge dense, arginine-rich protamine (38). (Protamines replace histones to achieve a 10-fold greater compaction of DNA in sperm.) Titration of salmon protamine into 16bp dsDNA gave a fully exothermic isotherm (Fig. 5E), thereby confirming the trend.

### Mutation of proline leads to structural changes

While the higher effectiveness of arginine over lysine in DNA compaction has been noted before, e.g. in studies of synthetic oligoarginine (39), the powerful DNA-condensing properties of CH1_PA_ were unexpected and its physical origin less clear. Proline is the most abundant amino acid in the H1.11L C-tail (13%) after lysine and alanine. It is also relatively well distributed: the *κ* parameter for proline alone is 0.134, and the two longest proline-free regions are 11 and 13 residues (Fig. 3). Proline has many known impacts on secondary structure: location at the ends of helices (40), ability to isomerise (41) and potential to form polyproline II helices (42). The structures of all the proteins were explored using circular dichroism (CD) spectroscopy.

The far-UV CD spectra for CH5, CH5_K_ and CH1_R_ were identical to CH1 and consistent with proteins that are predominantly disordered, having no significant features beyond the negative peak at ∼ 200 nm (Fig. 6A). Therefore, arginine content and charge density do not noticeably impact the structure of the free CH1 protein, as has been observed previously for CH1 vs CH5 (43). The reduced hydrophobicity mutants CH1_VA_ and CH1_VT_ were also indistinguishable from CH1 (Fig. 6B). However, CH1_PA_ showed a shallow dip around 222 nm and a shift of the ∼ 200 nm peak to higher wavelengths, indicative of a small percentage of alpha helix in a disordered background. On cooling from 25 °C to 0 °C, the amount of alpha helix visibly increased (Fig. 6C). This was in contrast to CH1 at 0 °C, which showed a smaller change and in the opposite direction: the peak ∼ 200 nm became more negative and there was a slight increase in the signal around 218 nm, indicating temperature-dependent stabilisation of polyproline II (PPII) helix, as expected for a lysine-rich polypeptide at low/neutral pH (44, 45).

**Figure 6.**
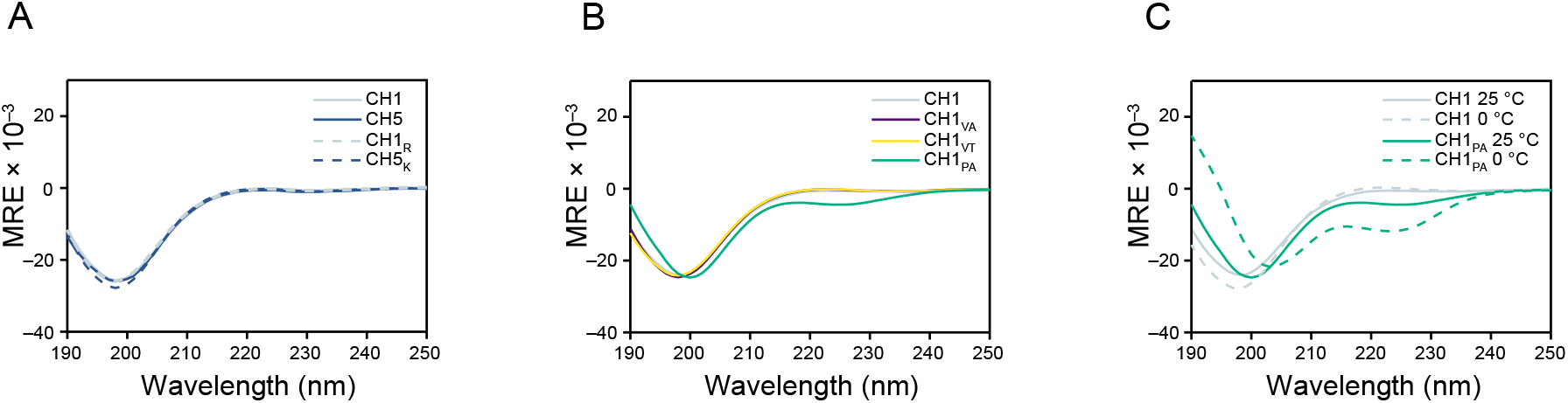
Far-UV CD of the free proteins. (A) Arginine and charge density series. (B) Hydrophobicity and proline knockouts. (C) Proline knockout and CH1 vs. temperature. Temperature 25 °C unless indicated, in buffer containing 150 mM NaF (total *I* = 160 mM).

In order to investigate this further, ^15^N-labelled CH1 and CH1_PA_ were expressed and purified. The ^15^N-HSQC spectrum of CH1 (Fig. 7A) displayed the limited ^1^H^N^ chemical shift dispersion and narrow line widths characteristic of a highly dynamic polypeptide chain, as observed previously (16), with a small degree of line broadening on cooling to 0 °C, consistent with a degree of stiffening of the chain due to PPII. The ^15^N-HSQC spectrum of CH1_PA_ at 25 °C was also consistent with a disordered protein (Fig. 7B), albeit with slightly increased ^1^H^N^ chemical shift dispersion and line widths, and a general shift upfield (lower ppm). However, the spectrum at 0 °C revealed a large number of peaks at non-random-coil positions, and significantly increased line widths, consistent with the secondary structure formation seen by CD (Fig. 6C).

**Figure 7.**
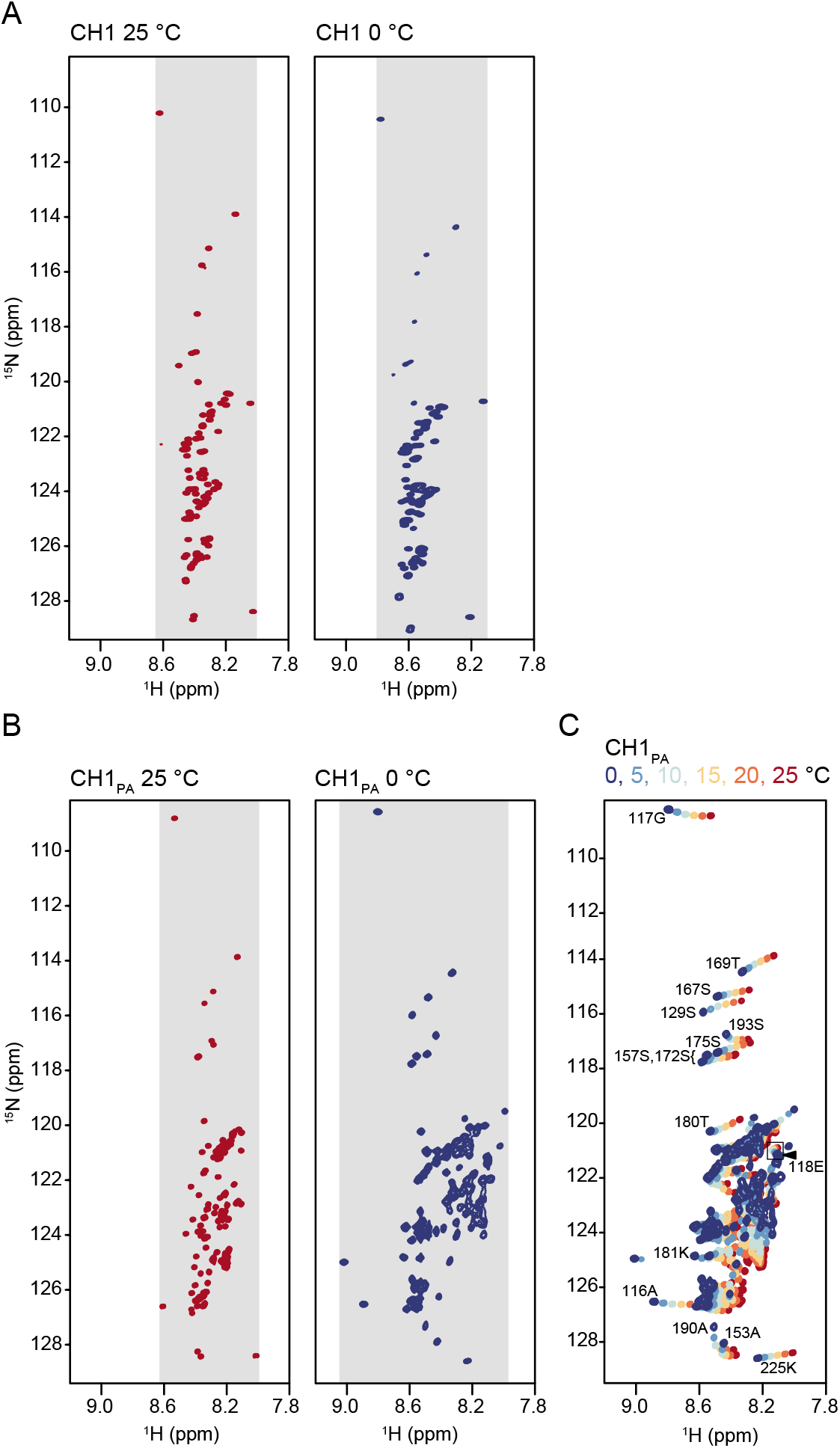
^15^N-HSQC NMR of the free proteins. (A) CH1 at 25 °C and 0 °C. (B) Proline knockout at 25 °C and 0 °C. (C) Proline knockout vs. temperature, with peak assignments. Grey shading present to illustrate chemical shift dispersion in ^1^H^N^. Buffer contains 150 mM NaCl (total *I* = 160 mM).

The temperature dependence of ^1^H^N^ chemical shifts is generally linear, and the rate is a reliable indicator of the extent of hydrogen bonding. Values of Δδ/ΔT less negative than –4.5 ppb/K are indicative of a stable hydrogen bond (46). The values for CH1, where the previously assigned peaks (16) can be reliably tracked, lie around –10 ppb/K, (mean = –9.9, σ = 0.9) with no single value less negative than –6.5 ppb/K (for Glu118). The values for CH1_PA_ were significantly less negative (e.g. Glu118 was –1.7 ppb/K, shown boxed in Fig. 7C). Furthermore, for CH1_PA_, many peaks (including Glu118) showed curved rather than straight trajectories of chemical shift with temperature, and peak broadening, consistent with the onset of folding. Peak assignment was made challenging by the ultra-low sequence complexity and repeated motifs (CH1_PA_ is 40% lysine, 43% alanine), but several peaks could nevertheless be assigned sequence-specifically, some having linear temperature trajectories (e.g. Ser175) and some curved (e.g. Ala153).

Next, we investigated the structures formed in the condensates by CD. Previously, for CH1, we observed a large negative ellipticity at ∼ 280 nm (16), which we inferred to be the result of scattering from a cholesteric liquid crystalline phase in which the DNA duplexes adopted a left-handed twisted stack (47), but only in low ionic strength conditions. Such signals have historically been denoted ‘polymer and salt induced-’, ‘psi-’, or ‘ψ’-DNA. CD spectra for the CH1, CH5, CH5_K_ and CH1_R_ droplet phases at physiological ionic strength are shown in Fig. 8A. No ψ-DNA signal was seen for CH1, consistent with our previous observations at physiological ionic strength (16), but the signal for CH5 is very different, showing a negative ellipticity at 295 nm consistent with ψ-DNA. The signals from CH5_K_ and CH1_R_ are in between, although CH1_R_ closely resembles CH1 while CH5_K_ deviates further from CH1 in the direction of CH5. The signal from the CH1_PA_-DNA condensate (Fig. 8B) is very different, showing a pronounced ψ-DNA scattering signal at 283 nm, consistent with higher-order structure formation with the highest degree of dichroic scattering. The secondary dip at 226 nm may indicate alpha helix formation in the protein, but is inconclusive due to the large background signal from DNA.

**Figure 8.**
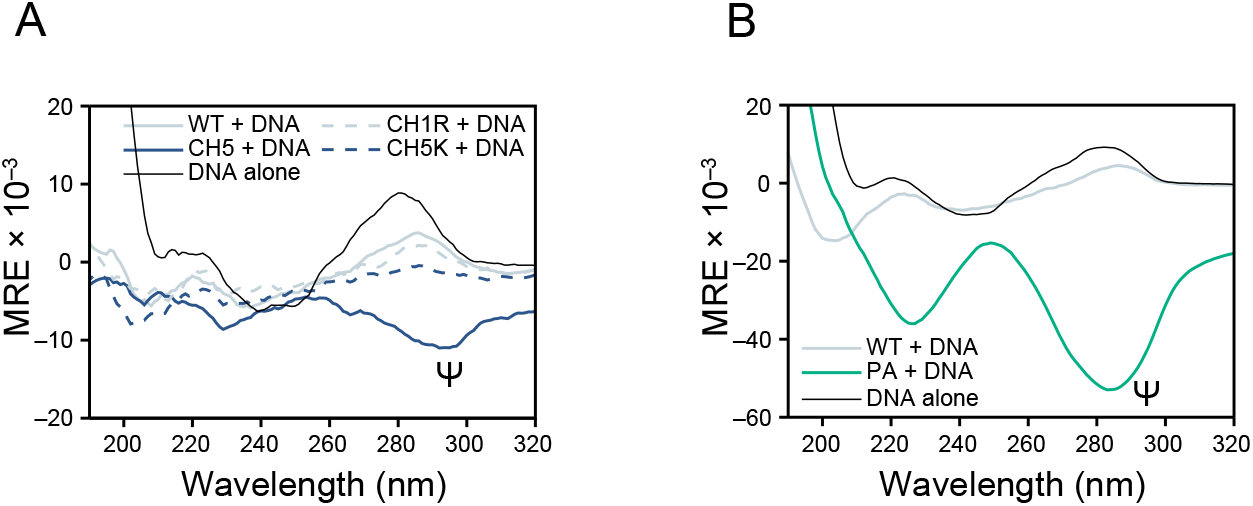
Far-UV CD of the protein:DNA complexes. (A) Arginine and charge density series and (B) CH1 and proline knockouts, with 20bp DNA, 1:1, at 25 °C in buffer containing 150 mM NaCl (total *I* = 160 mM). Free DNA (black) is also shown for reference. Temperature 25 °C unless indicated. The position of the ψ-DNA scattering signal is indicated (see the text).

## Discussion

The comparison of the lysine-rich CH1 and the arginine-rich and charge-dense CH5 was facilitated using a pair of carefully designed mutants. As expected, CH5 produces more robust condensation than CH1 (Fig. 4A & 5A). Dissecting this further, we find from the direct pairwise comparisons of CH1/CH1_R_ and CH5_K_/CH5 (which differ only in that 13 K are mutated to R) that high arginine content alone promotes condensation (Fig. 4A & 5B), as expected from its bidentate nature (21) and stacking propensity (24). A similar trend is seen for increasing positive charge density (compare CH1/CH5_K_, and CH1_R_/CH5); raising the fraction of positively charged residues from 0.4 to 0.5 promotes coacervation. However, we find that both high charge *and* arginine are needed for significant higher-order structure formation, or ψ-DNA, at physiological ionic strength (Fig. 8A). The underlying thermodynamics reveal a trend towards enthalpic control for the arginine-rich/charge dense linker histones relative to canonical H1s that are under a higher degree of entropic control (Fig. 5B).

Less expected were the effects of hydrophobic residues and proline. Mutation of all seven branched hydrophobic residues (valine) to alanine or threonine has a negligible effect on condensation (Fig. 4B & 5D, Supp. Fig. 3), as does 1,6- hexanediol, leading us to conclude that CH1/DNA condensation is a purely electrostatically driven process. However, the impact of proline on the robustness, thermodynamics and stoichiometry of condensation (Fig. 4C; Fig. 5C&D), and higher-order structuring (Fig. 8B) is profound. The reasons for this appear to lie in the accessible structures (Fig. 6 & 7): proline is both rigid and lacks an amide proton to stabilise alpha helices beyond the first few residues. Its absence can therefore confer a propensity for alpha helix, which has highly contrasting properties to the PPII helix (Fig. 9). An ideal PPII helix has backbone dihedral angles φ = −75° and ψ = +145°, resulting in a left-handed helix of precisely three residues per turn. The structure is extended (3.1 Å/residue) and without backbone hydrogen bonds, forming due to the repulsions between like-charged side-chains (in this case lysine), and is thought to be populated by many disordered proteins at least to some extent (44, 45). In contrast, an alpha helix is right-handed, hydrogen bonded and 3.6 residues per turn. Compared to the PPII helix, the structure is relatively compressed (1.5 vs 3.1 Å/residue), and side-chain repulsions are instead minimised through the stagger achieved by the 100° rotation per residue. These differences will directly impact the charge density of the protein, and also its ability to compress further on binding DNA. An AlphaFold prediction (48) shows CH1_PA_ to be mostly alpha helix, with a high confidence score, in contrast to CH1 (Supp. Fig. 4). By contrast, in our experiments, free CH1_PA_ is not fully alpha helical; even at 0 °C it is clear that formation of a complete and stable helix is not possible (Fig. 6C), presumably due to repulsions between lysine side chains that are too great even with the stagger. However, AlphaFold often predicts conditional folding (49), and CH1_PA_ is likely to compress further on binding DNA as the phosphate backbone reduces the repulsions, and indeed there are possible indications of alpha helix stabilisation in the CD spectra of the droplet phase (Fig. 8B). CH1 has 111 residues, and would therefore stretch to 34 nm if 100% PPII, but compress to 17 nm if fully alpha helical, some way closer to the length of dsDNA with the same overall charge (20bp, 7 nm). A closer match in charge density and rigidity to DNA is likely to be highly advantageous for binding and further condensation, and the three-dimensional nature of a helix could provide a more favourable template for higher-order structure formation. We propose that the ability to regulate charge density by compression in this spring-like manner underlies the two main differences that we observe between DNA condensation by CH1 and CH1_PA_: it becomes enthalpy driven, and occurs at lower protein:DNA stoichiometries, i.e. CH1_PA_ is more effective at charge-driven condensation, despite it having the same formal net charge and net charge per residue as CH1.

**Figure 9.**
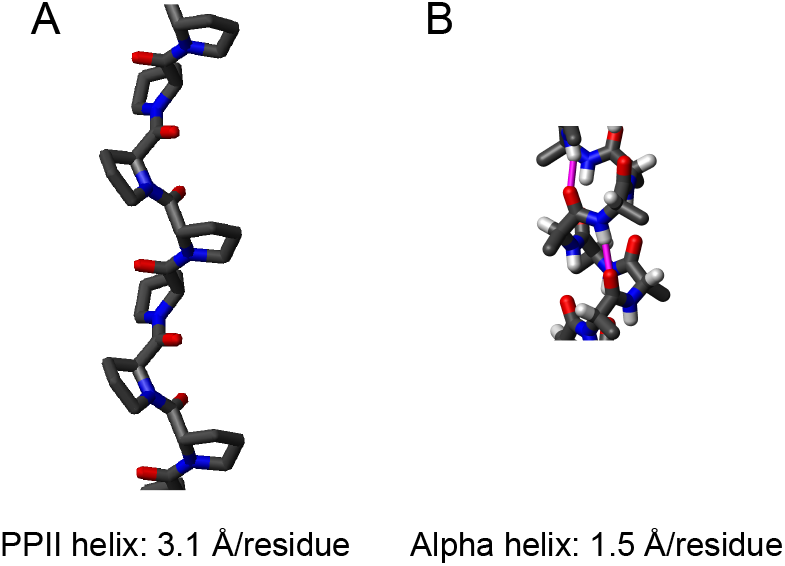
Idealised helical structures. Close-up side views of stick models of (A) polyproline II helix (PPII), and (B) alpha-helix. Original images created with MOLMOL (71) and made available under GFDL licence.

### Long proline-free regions are not uncommon in linker histone tails

Clues as to the possible biological relevance of proline across CH1s can be found by inspection of our curated database. Fig. 10 shows the length and incidence of proline-free regions in each CH1 across the set, including the three known avian CH5s, alongside NCPR and fraction arginine. Most proline-free regions are short (shown by the clustering of points along the bottom of the graph at ≤ 20 residues), as for H1.11L. However, several H1s have longer regions, of up to 213 residues. As for CH1_PA_, these are generally predicted to be helical by AlphaFold, with varying confidence scores. (A notable exception is the H1 from the fungus *Ashbya* (#81 H1_ASHGO) that might be expected to be helical given the proline-free region of 37 residues, however, an unusually high valine content (0.13, mean is 0.05, σ = 0.015) may prevent helix prediction in the proline gap by AlphaFold due to its lower incidence in helices (50)). Closer inspection of the helix-containing CH1s and their annotations (where present) reveals that many are thought to have gene repressive functions due to their expression in sperm or the testis e.g. #78 H1FNT_RAT and #84 H1FNT_MOUSE, the testis-specific H1s in rats and mice essential for normal spermatogenesis and male fertility (gaps 213 and 189, respectively, predicted helical). Helix is also predicted for #14 H1_PARAN, the H1 from the sperm of *Parechinus angulosus*, a sea urchin, which seems likely given the experimental observation of helix in the closely-related *Echinus esculentus* (43). The *Parechinus* H1, as well as a 57-residue proline-free region that like CH1_PA_ is very alanine-rich (0.47, not shown), also contains a moderate level of arginine overall (0.08), which presumably acts together with the proline-free region to confer more robust condensation leading to a gene-repressive function that must be resilient to the ionic conditions found in the sea to protect the paternal DNA. However it is by no means the case the drivers of condensation established here (arginine, positive charge density by net charge per residue, or by proline-free regions linked to helix formation) are all three present in gene-repressive H1s, e.g. the testis-specific #82 H1FNT_HUMAN has high arginine (0.24), a fairly long proline-free region (33 residues, predicted helical), but a relatively low charge density (NCPR 0.24). While it could be the case that arginine, NCPR and proline-free regions are three alternative ways for an organism to achieve the same outcome evolutionarily, it could also be the case that they are differently regulatable, conferring situation-specific advantages. In support of this, proline is required for the cyclin-dependent kinases that drive cell-cycle-dependent decondensation by phosphorylation (consensus site [S/T]<u>P</u>x[K/R]), so a proline-free region that is inert to this particular mechanism may have a functional niche in a cell-cycling background, a concept not unlike the recent report of ‘lysine deserts’ that prevent adventitious ubiquitylation (51).

**Figure 10.**
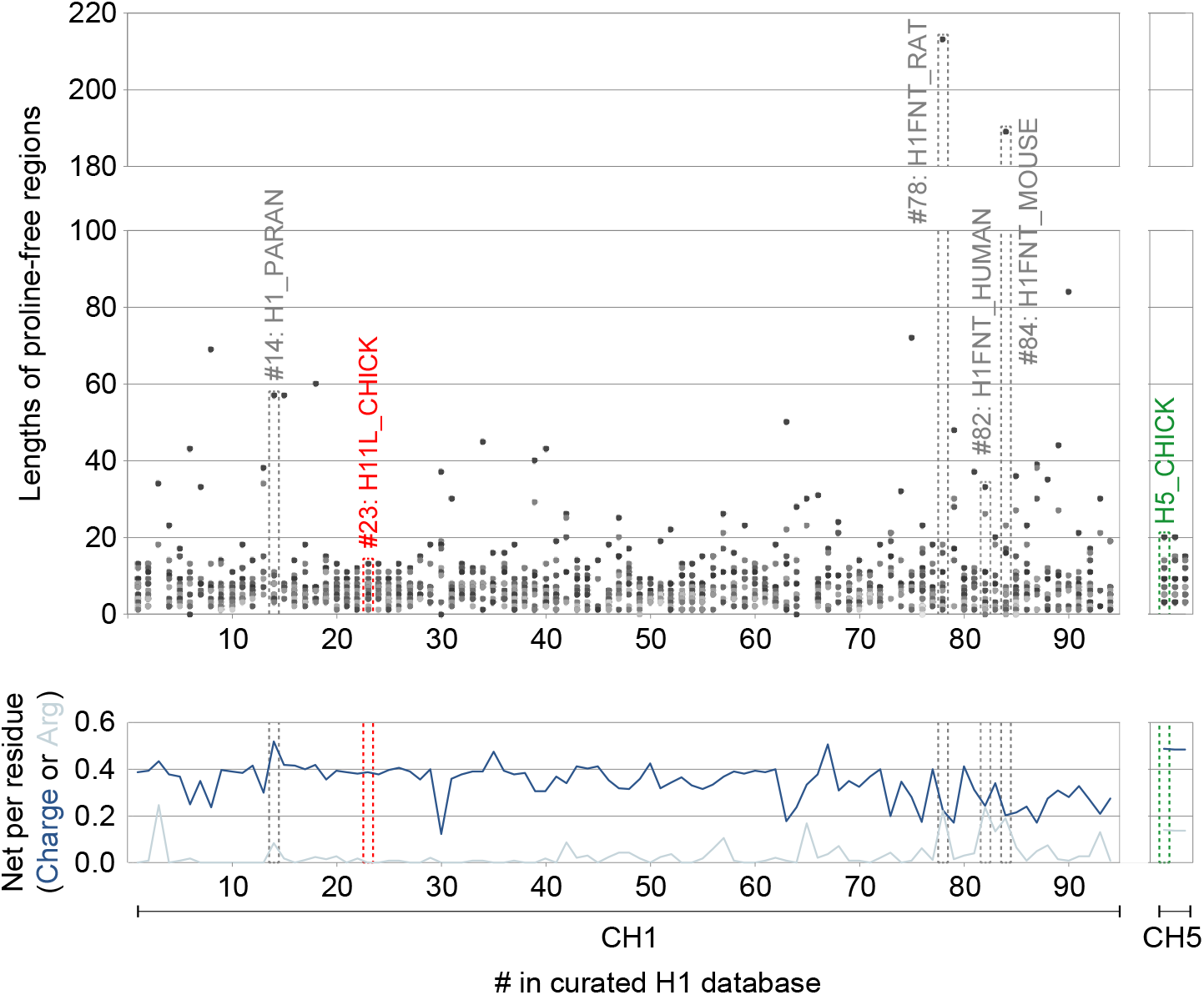
Proline-free regions across linker histones. (*Top*): All proline-free regions across the curated set of 94 CH1s are shown as dots corresponding to their length in residues, vs. record number in the database (Supp. Fig. S1). The three known avian CH5s are also shown on the right. (*Bottom*): Net charge per residue, and net arginine per residue (fraction arginine). Proteins with long proline-free regions and known gene-repressive functions (see text) are boxed in grey and labelled. CH1 and CH5 are boxed and labelled in red and green, respectively.

### Physiological relevance of the model system

Liquid-like behaviour in chromatin depends on the length scale of the measurement, a topic that has been recently and thoroughly reviewed (52). It follows that the length of the DNA also impacts the observed behaviour (53, 54): *In vivo*, long genomic DNA may form a constricted elastic gel scaffold on the micron scale that supports condensation of chromatin binding proteins (55). However, these structures, while globally constrained, retain high levels of local dynamics (10, 56). A question that naturally arises is whether *in vitro* studies with short DNA can be extrapolated to understand the local (nm-scale) properties of full-length chromatin. Shorter lengths of DNA (such as short oligos or *in vitro* 12-nucleosome arrays) are less elastically constrained and undergo phase separation into discrete μm-scale droplets (16, 20, 54, 57, 58). The same is true of fragmented chromatin: a recent study showed that nuclease-driven fragmentation of mitotic chromosomes resulted in droplets, but did not change the chromatin density, compaction state and resistance to perforation by microtubules (59). Taken together, these results demonstrate that the global physical constraints imposed by long DNA scaffolds do not necessarily define the local physicochemical properties, given these can be preserved when short oligos are released and become able to form droplets due to their relative translational and rotational freedom. The droplet state is therefore diagnostic of a condensation propensity, and its physicochemical properties can report on the local properties of chromatin. Historically, the use of *in vitro* models such as 12-nucleosome arrays has been justified on the basis that they show the same morphology (60), SAXS profiles and DNA accessibility (61) as endogenous chromatin. Our minimal model excludes nucleosomes, but the free energy contribution that nucleosome-nucleosome interactions make to condensation is relatively weak (∼–1.6 kcal/mol) (62) compared to CH1/DNA interactions, which are at least 5x greater, and often orders of magnitude more, depending on the context (16, 63, 64). Overall, our findings support the utility of *in vitro* models with short DNAs as they can reproduce the material properties of *in vivo* chromatin at short length scales.

## Conclusion

We have established a thermodynamic and structural basis for the range of DNA condensing properties of linker histones. In particular, we can rationalise the behaviour of those linked to gene-repressive functions in terminally differentiated cell types such as avian erythrocytes, and in the generation of sperm. We find that the predicted impact of high charge density and the replacement of lysine by arginine are both underpinned by enthalpic contributions that promote binding and condensation to a more ordered state. Further, we find a profound effect is seen for linker histones containing long proline-free regions, which is likely due to their higher achievable charge density though compact alpha-helical structures. Proline-free regions therefore provide an alternative thermodynamic switch to regulate condensation, structurally distinct from those with high charge density and arginine, and may offer further distinct properties in their regulation due to their immunity to the action of cyclin-dependent kinases.

## Materials and Methods

### Protein expression and purification

Codon optimised sequences for the various CH1s were obtained by gene synthesis. Proteins were expressed and purified as described previously for CH1 (16), although the yields were severely reduced for several mutants, and both CH5 and CH1_PA_ required higher salt to elute from the 5 mL HiTrap SP HP column. Salmon protamine was purchased from Sigma. Oligonucleotides were purchased from Merck or IDT, and annealed as described previously (16).

### Concentration measurements

Protein and DNA concentrations were determined spectrophotometrically using a NanoDrop One^C^ instrument (ThermoFisher). A_205_ was used for the H1 constructs due to their low ε_280_. Extinction coefficients were calculated using http://nickanthis.com/tools/a205.html based on Anthis & Clore, 2013 (65). Turbidity was measured in 50 or 100 μl, 1 cm path length microcuvettes by A_340_.

### Isothermal Titration Calorimetry

Experiments were performed on an ITC200 (GE/Malvern) at 25 °C. Samples were dialysed extensively into 10 mM sodium phosphate pH 6, 150 mM NaCl. The protein (at 100 μM for CH1s and 225 μM for protamine) was injected into the DNA (at 10 μM); 18 injections of 2 μl protein were performed, at intervals of 120 or 150 s with stirring at 750 rpm. Baseline correction and integration were performed in Origin.

### Far-UV Circular Dichroism Spectroscopy

Experiments on the free proteins were performed at ∼0.1 mg/ml over a 190-250 nm range at the indicated temperature in 10 mM sodium phosphate pH 6, 150 mM NaF, and in 1 mm path-length cuvettes. Spectra were acquired using an AVIV 410 spectrometer in 1 nm wavelength steps, averaged over three accumulations and baseline-corrected using buffer before smoothing, using the manufacturer’s software. Millidegree units were converted to mean residue ellipticity (MRE) with units deg cm^2^ dmol^-1^ res^-1^ using MRE = millideg. / {(no. residues – 1) × c × *l* × 10}, where c = molar concentration and *l* = path length in cm. For experiments on the protein:DNA 1:1 complexes, the wavelength range was extended to 320 nm, and a single scan was acquired per sample, to minimise any inconsistencies from droplets settling over time.

### Nuclear Magnetic Resonance

NMR measurements were made on ^15^N-labelled proteins (50-100 μM) in 10 mM sodium phosphate pH 6, 150 mM NaCl (NMR buffer) and 10-15% ^2^H_2_O. Experiments were recorded at the temperature indicated on a Bruker DRX800 spectrometer. Where possible, assignments were obtained using a conventional triple-resonance approach (HNCA, HN(CO)CA, HNCO, HNCACB, HN(CO)CACB) alongside TOCSY-^15^N-HSQC, NOESY-^15^N-HSQC and HNN/HN(C)N experiments (66, 67). Data were processed using AZARA (v.2.8, © 1993-2023; Wayne Boucher and Department of Biochemistry, University of Cambridge). Triple resonance experiments were recorded with 25% nonuniform sampling, using Poisson-gap sampling (68), and reconstructed using the Cambridge CS package and the CS-IHT algorithm (69). Assignments were made using CcpNmr Analysis v. 2.4 (70). Chemical shifts were referenced to 2,2-dimethyl-2-silapentane-5-sulfonic acid (DSS).

## Acknowledgements

We thank Jeremy Schmit for helpful discussions, and the NMR and Biophysics Facilities in the Department of Biochemistry, University of Cambridge for access to instrumentation. This work was supported by the Biotechnology and Biological Sciences Research Council (BB/T015403/1) and the Erasmus programme (studentship to E.D.).

## References

1. Gilbert N, Allan J (2019) The many length scales of DNA packaging. Essays Biochem 63:13–16.

2. Kornberg RD, Lorch Y (1999) Twenty-five years of the nucleosome, fundamental particle of the eukaryote chromosome. Cell 98(3):285–294.

3. Bednar J, Hamiche A, Dimitrov S (2016) H1-nucleosome interactions and their functional implications. Biochim Biophys Acta - Gene Regul Mech 1859(3):436– 443.

4. Thoma F, Koller TH, Klug A (1979) Involvement of histone H1 in the organization of the nucleosome and of the salt-dependent superstructures of chromatin. J Cell Biol 83(November):403–427.

5. Woodcock CL (1994) Chromatin fibers observed in situ in frozen hydrated sections. Native fiber diameter is not correlated with nucleosome repeat length. J Cell Biol 125(1):11–19.

6. Scheffer MP, Eltsov M, Frangakis AS (2011) Evidence for short-range helical order in the 30-nm chromatin fibers of erythrocyte nuclei. Proc Natl Acad Sci U S A 108(41):16992–16997.

7. Hou Z, Nightingale F, Zhu Y, MacGregor-Chatwin C, Zhang P (2023) Structure of native chromatin fibres revealed by Cryo-ET in situ. Nat Commun 14:6324(October):2023.09.03.556082.

8. Maeshima K, Ide S, Hibino K, Sasai M (2016) Liquid-like behavior of chromatin. Curr Opin Genet Dev 37:36–45.

9. Maeshima K, Hihara S, Eltsov M (2010) Chromatin structure: Does the 30-nm fibre exist in vivo? Curr Opin Cell Biol 22(3):291–297.

10. Nozaki T, et al. (2017) Dynamic Organization of Chromatin Domains Revealed by Super-Resolution Live-Cell Imaging. Mol Cell 67(2):282–293.e7.

11. Hihara S, et al. (2012) Local Nucleosome Dynamics Facilitate Chromatin Accessibility in Living Mammalian Cells. Cell Rep 2(6):1645–1656.

12. Allan J, Hartman PG, Crane-Robinson C, Aviles FX (1980) The structure of histone H1 and its location in chromatin. Nature 288(18):675–679.

13. Syed S, et al. (2010) Single-base resolution mapping of H1–nucleosome interactions and 3D organization of the nucleosome. Proc Natl Acad Sci U S A 107(21):9620–9625.

14. Bednar J, et al. (2017) Structure and Dynamics of a 197 bp Nucleosome in Complex with Linker Histone H1. Mol Cell 66(3):384–397.e8.

15. Meyer S, et al. (2011) From crystal and NMR structures, footprints and cryo-electron-micrographs to large and soft structures: nanoscale modeling of the nucleosomal stem. Nucleic Acids Res 39(21):9139–54.

16. Turner AL, et al. (2018) Highly disordered histone H1-DNA model complexes and their condensates. Proc Natl Acad Sci U S A 115(47):11964–11969.

17. Watson M, Stott K (2019) Disordered domains in chromatin-binding proteins. Essays Biochem 63(1):147–156.

18. Hendzel MJ, Lever MA, Crawford E, Th’Ng JPH (2004) The C-terminal Domain Is the Primary Determinant of Histone H1 Binding to Chromatin in Vivo. J Biol Chem 279(19):20028–20034.

19. Allan J, Mitchell T, Harborne N, Bohm L, Crane-Robinson C (1986) Roles of H1 domains in determining higher order chromatin structure and H1 location. J Mol Biol 187(4):591–601.

20. Gibson BA, et al. (2019) Organization of Chromatin by Intrinsic and Regulated Phase Separation. Cell 179:1–15.

21. Ichimura S, Mita K, Zama M (1978) Conformation of poly(L-arginine). I. Effects of anions. Biopolymers 17(12):2769–2782.

22. Derouchey J, Hoover B, Rau DC (2013) A comparison of DNA compaction by arginine and lysine peptides: A physical basis for arginine rich protamines. Biochemistry 52(17):3000–3009.

23. Vondrášek J, Mason PE, Heyda J, Collins KD, Jungwirth P (2009) The molecular origin of like-charge arginine - Arginine pairing in water. J Phys Chem B 113(27):9041–9045.

24. Neves MAC, Yeager M, Abagyan R (2012) Unusual arginine formations in protein function and assembly: Rings, strings, and stacks. J Phys Chem B 116(23):7006–7013.

25. Izzo A, Kamieniarz K, Schneider R (2008) The histone H1 family: Specific members, specific functions? Biol Chem 389(4):333–343.

26. Hergeth SP, Schneider R (2015) The H1 linker histones: multifunctional proteins beyond the nucleosomal core particle. EMBO Rep 16(11):1439–1453.

27. Millán-Ariño L, Izquierdo-Bouldstridge A, Jordan A (2016) Specificities and genomic distribution of somatic mammalian histone H1 subtypes. Biochim Biophys Acta - Gene Regul Mech 1859(3):510–519.

28. Neelin JM, Callahan PX, Lamb DC, Murray K (1964) The histones of chicken erythrocyte nuclei. Can J Biochem 42(12):1743–1752.

29. Zhou BR, et al. (2016) A Small Number of Residues Can Determine if Linker Histones Are Bound On or Off Dyad in the Chromatosome. J Mol Biol 428(20):3948–3959.

30. Zhou BR, et al. (2021) Distinct Structures and Dynamics of Chromatosomes with Different Human Linker Histone Isoforms. Mol Cell 81(1):166–182.e6.

31. Andrés M, García-Gomis D, Ponte I, Suau P, Roque A (2020) Histone H1 Post-Translational Modifications: Update and Future Perspectives. Int J Mol Sci 21:1–22.

32. Das RK, Pappu R V. (2013) Conformations of intrinsically disordered proteins are influenced by linear sequence distributions of oppositely charged residues. Proc Natl Acad Sci 110(33):13392–13397.

33. Holehouse AS, Das RK, Ahad JN, Richardson MOG, Pappu R V. (2017) CIDER: Resources to Analyze Sequence-Ensemble Relationships of Intrinsically Disordered Proteins. Biophys J 112(1):16–21.

34. Cato L, Stott K, Watson M, Thomas JO (2008) The interaction of HMGB1 and linker histones occurs through their acidic and basic tails. J Mol Biol 384(5):1262–72.

35. Kroschwald S, Maharana S, Alberti S (2017) Hexanediol: a chemical probe to investigate the material properties of membrane-less compartments. Matters:1–6.

36. Mascotti DP, Lohman TM (1990) Thermodynamic extent of counterion release upon binding oligolysines to single-stranded nucleic acids. Proc Natl Acad Sci 87(8):3142–3146.

37. Vitorazi L, et al. (2014) Evidence of a two-step process and pathway dependency in the thermodynamics of poly(diallyldimethylammonium chloride)/poly(sodium acrylate) complexation. Soft Matter 10(i):9496–9505.

38. DeRouchey JE, Rau DC (2011) Role of amino acid insertions on intermolecular forces between arginine peptide condensed dna helices: Implications for protamine-DNA packaging in sperm. J Biol Chem 286(49):41985–41992.

39. Mascotti DP, Lohman TM (1997) Thermodynamics of oligoarginines binding to RNA and DNA. Biochemistry 36(23):7272–7279.

40. 40. Richardson JS, Richardson DC (1988) Amino Acid Preferences for Specific Locations at the Ends of <symbol>a</symbol> Helices. Science 240(11):1648–1652.

41. MacArthur MW, Thornton JM (1991) Influence of Proline Residues on Protein Conformation. J Mol Biol 218:397–412.

42. Tiffany ML, Krimm S (1968) Circular Dichroism of Poly-L-proline in an Unordered Conformation. Biopolymers 6:1767–1770.

43. Clark DJ, Hill CS, Martin SR, Thomas JO (1988) Alpha-helix in the carboxy-terminal domains of histones H1 and H5. EMBO J 7(1):69–75.

44. Tiffany ML, Krimm S (1968) New chain conformations of poly(glutamic acid) and polylysine. Biopolymers 6:1379–1382.

45. Rucker AL, Creamer TP (2002) Polyproline II helical structure in protein unfolded states: Lysine peptides revisited. Protein Sci 11(4):980–985.

46. Baxter NJ, Williamson MP (1997) Temperature dependence of 1H chemical shifts in proteins. J Biomol NMR 9(4):359–369.

47. Maestre MF, Reich C (1980) Contribution of Light Scattering to the Circular Dichroism of Deoxyribonucleic Acid Films, Deoxyribonucleic Acid-Polylysine Complexes, and Deoxyribonucleic Acid Particles in Ethanolic Buffers. Biochemistry 19(23):5214–5223.

48. Mirdita M, et al. (2022) ColabFold: making protein folding accessible to all. Nat Methods 19(6):679–682.

49. Alderson TR, Pritišanac I, Kolarić E, Moses AM, Forman-Kay JD (2023) Systematic identification of conditionally folded intrinsically disordered regions by AlphaFold2. bioRxiv:2022.02.18.481080.

50. Pace CN, Scholtz JM (1998) A helix propensity scale based on experimental studies of peptides and proteins. Biophys J 75(1):422–427.

51. Kampmeyer C, et al. (2023) Lysine deserts prevent adventitious ubiquitylation of ubiquitin-proteasome components. Cell Mol Life Sci 80(6):1–18.

52. Hansen JC, Maeshima K, Hendzel MJ (2021) The solid and liquid states of chromatin. Epigenetics and Chromatin 14(1):1–33.

53. Muzzopappa F, Hertzog M, Erdel F (2021) DNA length tunes the fluidity of DNA-based condensates. Biophys J 120(7):1288–1300.

54. Chen Q, et al. (2022) Chromatin Liquid–Liquid Phase Separation (LLPS) Is Regulated by Ionic Conditions and Fiber Length. Cells 11(3145):1–14.

55. Strickfaden H, et al. (2020) Condensed Chromatin Behaves like a Solid on the Mesoscale In Vitro and in Living Cells. Cell 183(7):1772–1784.e13.

56. 56. Gibson BA, et al. (2021) In Diverse Conditions Intrinsic Chromatin Condensates Have Liquid-like Material Properties. bioRxiv:2021.11.22.469620.

57. Shakya A, King JT (2018) Non-Fickian Molecular Transport in Protein-DNA Droplets. ACS Macro Lett 7(10):1220–1225.

58. Shakya A, Park S, Rana N, King JT (2019) Liquid-Liquid Phase Separation of Histone Proteins in Cells: Role in Chromatin Organization. Biophys J 4:1–12.

59. Schneider MWG, et al. (2022) A mitotic chromatin phase transition prevents perforation by microtubules. Nature 609(7925):183–190.

60. Carruthers LM, Bednar J, Woodcock CL, Hansen JC (1998) Linker histones stabilize the intrinsic salt-dependent folding of nucleosomal arrays: Mechanistic ramifications for higher-order chromatin folding. Biochemistry 37(42):14776–14787.

61. Maeshima K, et al. (2016) Nucleosomal arrays self-assemble into supramolecular globular structures lacking 30-nm fibers. EMBO J 35(10):1115– 1132.

62. Funke JJ, et al. (2016) Uncovering the forces between nucleosomes using DNA origami. Sci Adv 2(11). doi:10.1126/sciadv.1600974.

63. Machha VR, et al. (2013) Calorimetric studies of the interactions of linker histone H10 and its carboxyl (H10-C) and globular (H10-G) domains with calf-thymus DNA. Biophys Chem 184:22–28.

64. White AE, Hieb AR, Luger K (2016) A quantitative investigation of linker histone interactions with nucleosomes and chromatin. Sci Rep 6:19122.

65. Anthis NJ, Clore GM (2013) Sequence-specific determination of protein and peptide concentrations by absorbance at 205 nm. Protein Sci 22(6):851–858.

66. Cavanagh J, Skelton NJ, Fairbrother WJ, Rance M, Palmer AGI (2006) Protein NMR Spectroscopy Principles and Practice (Academic Press). 2nd Ed.

67. Panchal SC, Bhavesh NS, Hosur R V (2001) Improved 3D triple resonance experiments, HNN and HN(C)N, for HN and 15N sequential correlations in (13C,15N) labeled proteins: Application to unfolded proteins. J Biomol NMR 20(C):135–147.

68. Hyberts SG, Takeuchi K, Wagner G (2010) Poisson-gap sampling and forward maximum entropy reconstruction for enhancing the resolution and sensitivity of protein NMR data. J Am Chem Soc 132(7):2145–2147.

69. Bostock MJ, Holland DJ, Nietlispach D (2012) Compressed sensing reconstruction of undersampled 3D NOESY spectra: Application to large membrane proteins. J Biomol NMR 54(1):15–32.

70. Vranken WF, et al. (2005) The CCPN data model for NMR spectroscopy: Development of a software pipeline. Proteins Struct Funct Genet 59(4):687– 696.

71. Koradi R, Billeter M, Wüthrich K (1996) MOLMOL: A program for display and analysis of macromolecular structures. J Mol Graph 14(1):51–55.

