## Supplementary Information for "A DNA condensation code for linker histones"

**A DNA condensation code for linker histones – *Supplementary information***

This PDF file includes Figures S1 to S4 and Supplementary References.

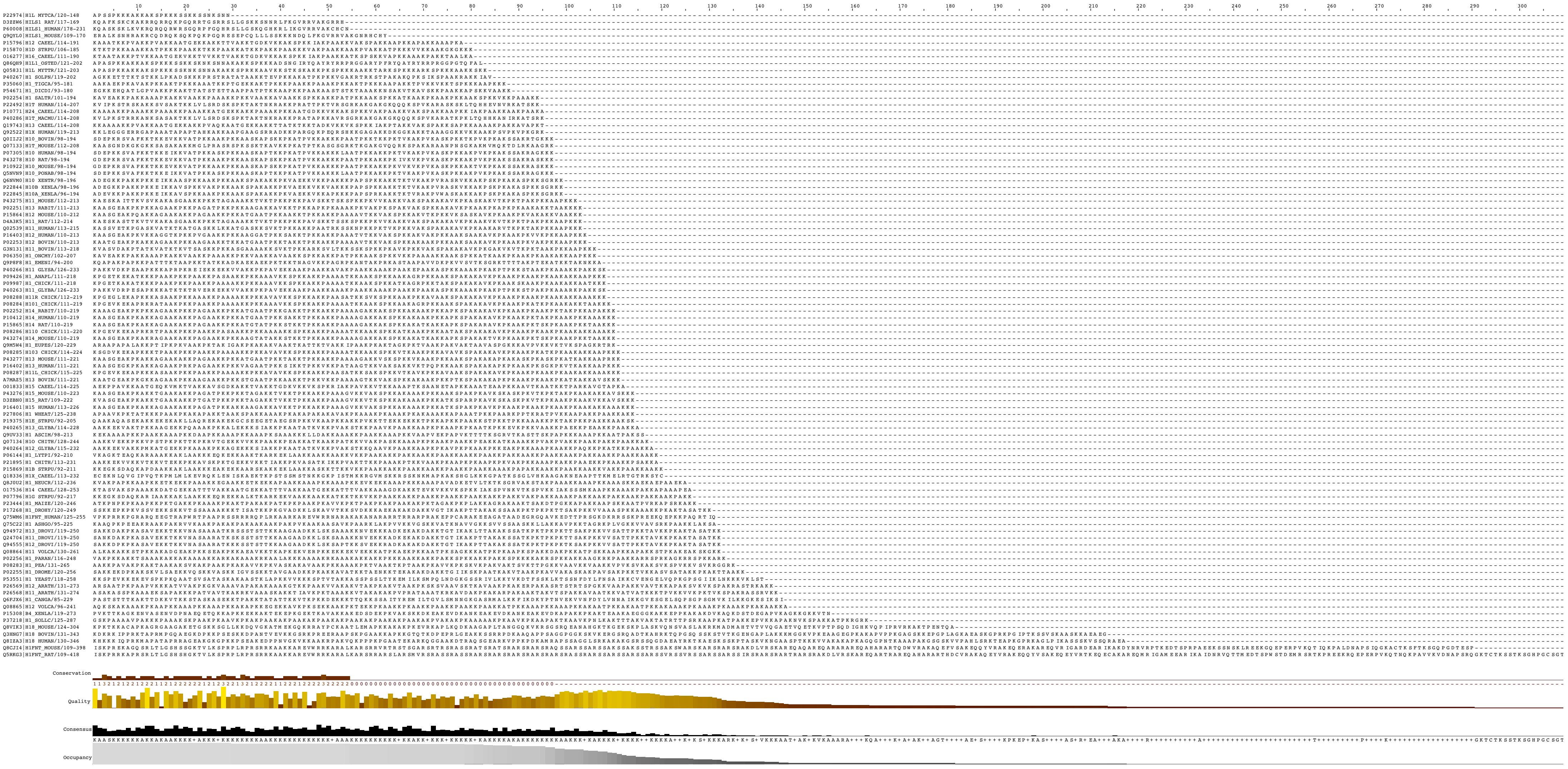

**Figure S1. Curated set of 94 H1 C-terminal tail sequences.** UniprotKB/SwissProt sequences extracted after alignment on the globular domains, displayed in order of increasing length. Figure prepared using Jalview v2 (1).

A Somatic linker histone H1 C-terminal tails in *Gallus gallus*

```
P08287|H11L_CHICK/115-225 KPGEVKEKAPKKKASAAKPKKPAAKKPAAAAKKPKKAVAVKKSPKKAKKPAASATKKSAKSPKKVTKAVKPKKAVAAKSPAKAKAVKPKAAKPKAAKPKAAKAKKAAAKKK
P08288|H11R_CHICK/112-219 KPGEGLEKAPKKKASAAKPKKAAAKKPAAAAKKPKKAVAVKKSPKKAKKPAASATKKSVKSPKK---AAKPKKAVAAKSPAKAKAVKPKAAKPKAAKPKAAKAKKAAAKKK
P08284|H101_CHICK/111-219 KPGEVKEKAPRKRATAAKPKKPAAKKPAAAAKKPKKAAAVKKSPKKAKKPAAAATKKAAKSPKKAAKAGRPKKA--AKSPAKAKAVKPKAAKPKATKPKAAKAKKTAAKKK
P08285|H103_CHICK/114-224 KSGDVKEKAPKKKTPAAKPKKPAAKKPAAAAKKPKKAVAVKKSPKKAKKPAAAATKKAAKSPKKVTKAAKPKKAVAVKSPAKAKAVKPKAAKPKATKPKAAKAKKAAPKKK
P09987|H1_CHICK/111-218   KPGETKAKATKK-KPAAKPKKPAAKKPAAAAKKPKKAAAVKKSPKKAKKPAAAATKKAAKSPKKATKAGRPKKT--AKSPAKAKAVKPKAAKSKAAKPKAAKAKKAATKKK
P08286|H110_CHICK/111-220 KPGEVKEKAPRKRTPAAKPKKPAAKKPASAAKKPKKAAAAKKSPKKAKKPAAAATKKAAKSPKKATKAAKPKKAATAKSPAKAKAVKPKAAKPKAAKPKAAKAKKAAAKK-
```

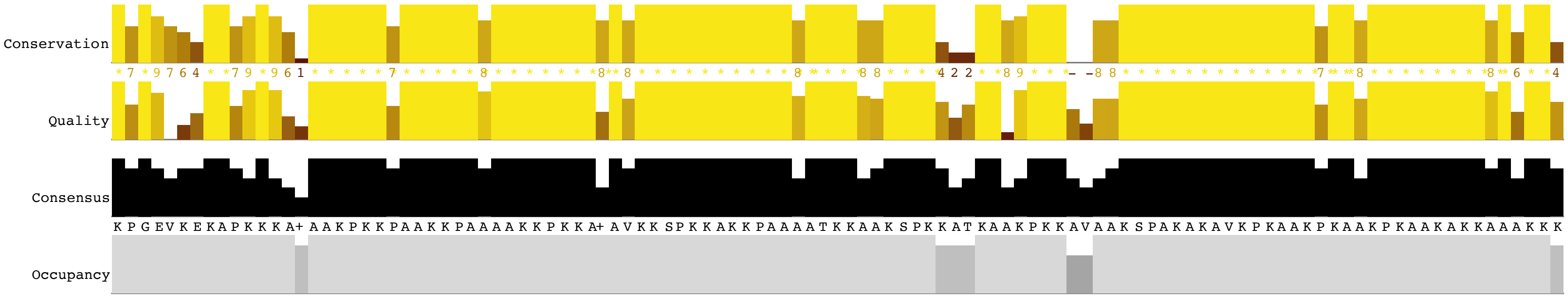


B Erythrocyte-specific linker histone H5 C-terminal tail in *Gallus gallus*

```
P02259|H5_CHICK/99-190    SDKAKRSPGKKKKAVRRSTSPKKAARPRKARSPAKKPKATARKARKKSRASPKKAKKPKTVKAKSRKASKAKKVKRSKPRAKSGARKSPKKK
```

**Figure S2. Linker histone C-terminal tails in *Gallus gallus*.** (A) Somatic H1 isoforms. (B) H5 variant. Arginine and proline are highlighted in dark blue and green, respectively. Branched hydrophobics (primarily valine) in yellow. Alignment and statistics prepared using Jalview v2 (1).

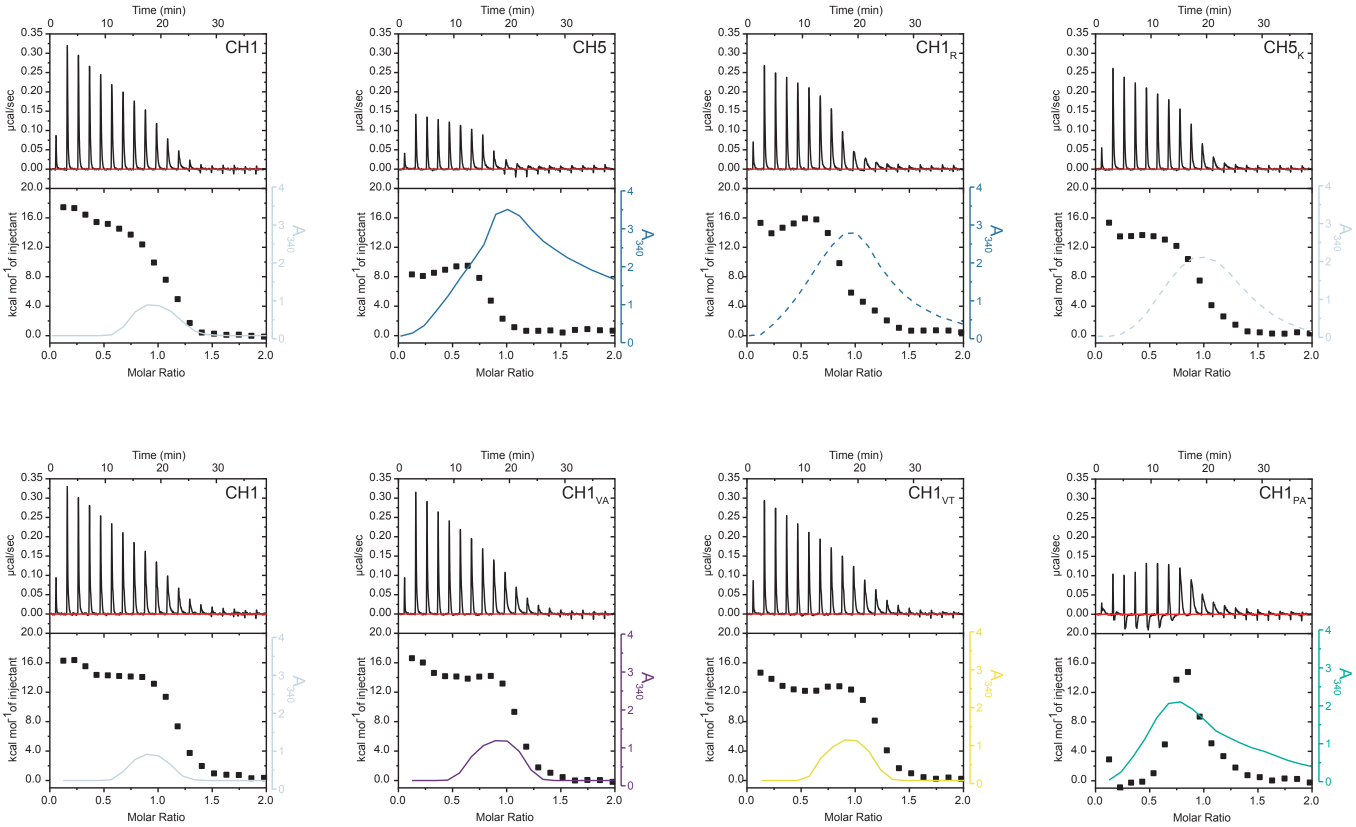


**Figure S3. Isothermal titration calorimetry (complete set).** Titrations of protein (as indicated) into 20 bp DNA, alongside $A_{340}$ to identify phase separation events. Raw data, isotherms and $A_{340}$ are shown, all recorded under the same conditions (25 °C, 10 mM sodium phosphate pH 6, 150 mM NaCl, protein concentration 100 µM, 20 bp DNA 10 µM).

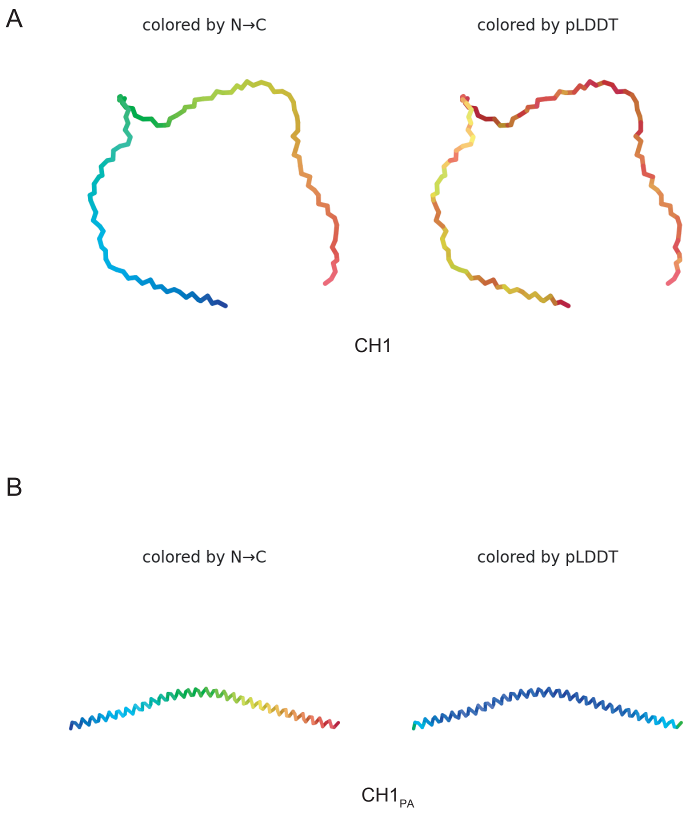


**Figure S4. AlphaFold2 predictions.** (A) CH1, (B) proline knockout, predicted using ColabFold (2). Confidence measure (pLDDT) colouring is blue (high confidence) to red (low confidence).

### Supplementary References

1. Waterhouse AM, Procter JB, Martin DMA, Clamp M, Barton GJ (2009) Jalview Version 2-A multiple sequence alignment editor and analysis workbench. *Bioinformatics* 25(9):1189–1191.

2. Mirdita M, et al. (2022) ColabFold: making protein folding accessible to all. *Nat Methods* 19(6):679–682.
